# Flow-driven delivery boosts antibiotic effectiveness by overwhelming bacterial defenses

**DOI:** 10.64898/2026.05.22.727216

**Authors:** Alexander M. Shuppara, Matthias D. Koch, Joseph E. Sanfilippo

## Abstract

Physical, chemical, and biological factors collectively determine antibiotic effectiveness. However, laboratory studies often consider only one or two of these factors and fail to capture key interaction effects. Here, we use microfluidics to study the interplay of host-relevant shear flow, the antibiotic gentamicin, and the human pathogen *Pseudomonas aeruginosa*. We discover bacterial populations can defend against gentamicin by chemical inactivation or physical sequestration. In low flow regimes, bacterial defenses succeed by eliminating gentamicin faster than it is delivered. As flow increases, delivery overwhelms bacterial defenses, allowing gentamicin to penetrate populations and inhibit cells. Combining biophysical simulations and long-channel microfluidics, we demonstrate that antibiotic susceptibility depends on spatial context, as cells at the beginning of the channel shield cells at the end. Surprisingly, we show that populations of sensitive cells can successfully defend against gentamicin, resulting in spatial gradients that are shaped by flow and cell density. Collectively, our interdisciplinary experiments reveal how flow overwhelms bacterial defenses, providing a framework to better understand antibiotic effectiveness in dynamic environments.

## Introduction

The human body must generate flow to survive. Bacterial pathogens must contend with host-generated flow, but they are typically studied in simplified conditions that lack flow. Recent microfluidic studies have revealed how flow can influence bacteria in unexpected ways (*1–3*). For example, flow can enhance adhesion (*4–8*), promote upstream motility (*9–15*), and shape biofilm architecture (*7*, *16–19*). In addition to applying a physical force, flow can alter chemical transport by washing away or replenishing small molecules. Flow-driven transport can promote growth in nutrient-limited regimes (*20–23*), potentiate the effects of chemical stress (*24–26*), and inhibit cell-cell communication (*27*, *28*). Although it is now clear that flow has widespread effects on bacterial survival, studies that focus on the interaction between bacterial pathogens and flow are still limited.

Antibiotic effectiveness is a key determinant of human health (*29*). As the list of effective antibiotics declines (*30*, *31*), new perspectives are needed. Bacteria have evolved several antibiotic defense strategies, including chemical inactivation and population-level shielding (*32*). In populations, cells on the periphery can protect cells on the interior by preventing delivery (*33*, *34*). Peripheral cells can prevent delivery in multiple ways: by chemically inactivating antibiotics or by physically blocking their access. Conceptually, flow could enhance antibiotic effectiveness by delivering antibiotics faster than bacteria can inactivate them (*35*). Similarly, flow could overcome the ability of bacterial populations to physically block access to all cells. As antibiotic delivery is often a rate-limiting step in treatment, there is an urgent need to understand the impact of flow on antibiotic delivery into bacterial populations.

Here, we use our microfluidic system to examine the interactions between host-relevant shear flow, the aminoglycoside gentamicin, and the human pathogen *Pseudomonas aeruginosa*. We quantitatively establish that gentamicin effectiveness is determined by the combinatorial effects of shear rate and population density. We show that the speed of gentamicin delivery determines effectiveness independent of the total dose delivered. To our surprise, we learned that populations of sensitive cells create spatial gradients, due to physical sequestration of gentamicin at the start of the channel. By joining multiple microfluidic devices, we discover that sensitive and resistant bacterial populations remove gentamicin at the same rate, highlighting the importance of physical sequestration. Our results demonstrate that flow promotes antibiotic effectiveness and incorporating flow has the potential to improve antibiotic discovery, diagnostic testing, and treatment strategies.

## Results

To understand how flow impacts antibiotic treatment of bacterial populations, we studied the interactions between flow, gentamicin, and *P. aeruginosa* cells. We precisely controlled flow by connecting a syringe pump to custom-fabricated microfluidic devices, which contained surface-attached cells (Figure 1A and S1). To study antibiotic resistance, we used a gentamicin-resistant *P. aeruginosa* strain that can inactivate gentamicin via acetylation (Figure S2). Based on our previous work (*35*), we hypothesized that higher flow could overcome gentamicin resistance. Flow can be described using shear rate, which is determined by flow rate and channel dimensions. To directly test our hypothesis, we delivered 100 µg/mL gentamicin into our microfluidic devices at various shear rates (0, 80, 160, 240, and 800 s^-1^). We chose these shear rates as they are commonly found in the human bloodstream (*36*), urinary tract (*37*), and lungs (*38*). Consistent with our hypothesis, no flow (0 s^-1^) and low flow (80 s^-1^) conditions resulted in robust bacterial growth, while higher shear rates (160, 240, 800 s^-1^) led to a stepwise decrease in growth (Figure 1B). Thus, higher flow can overcome *P. aeruginosa* resistance to gentamicin.

**Figure 1:**
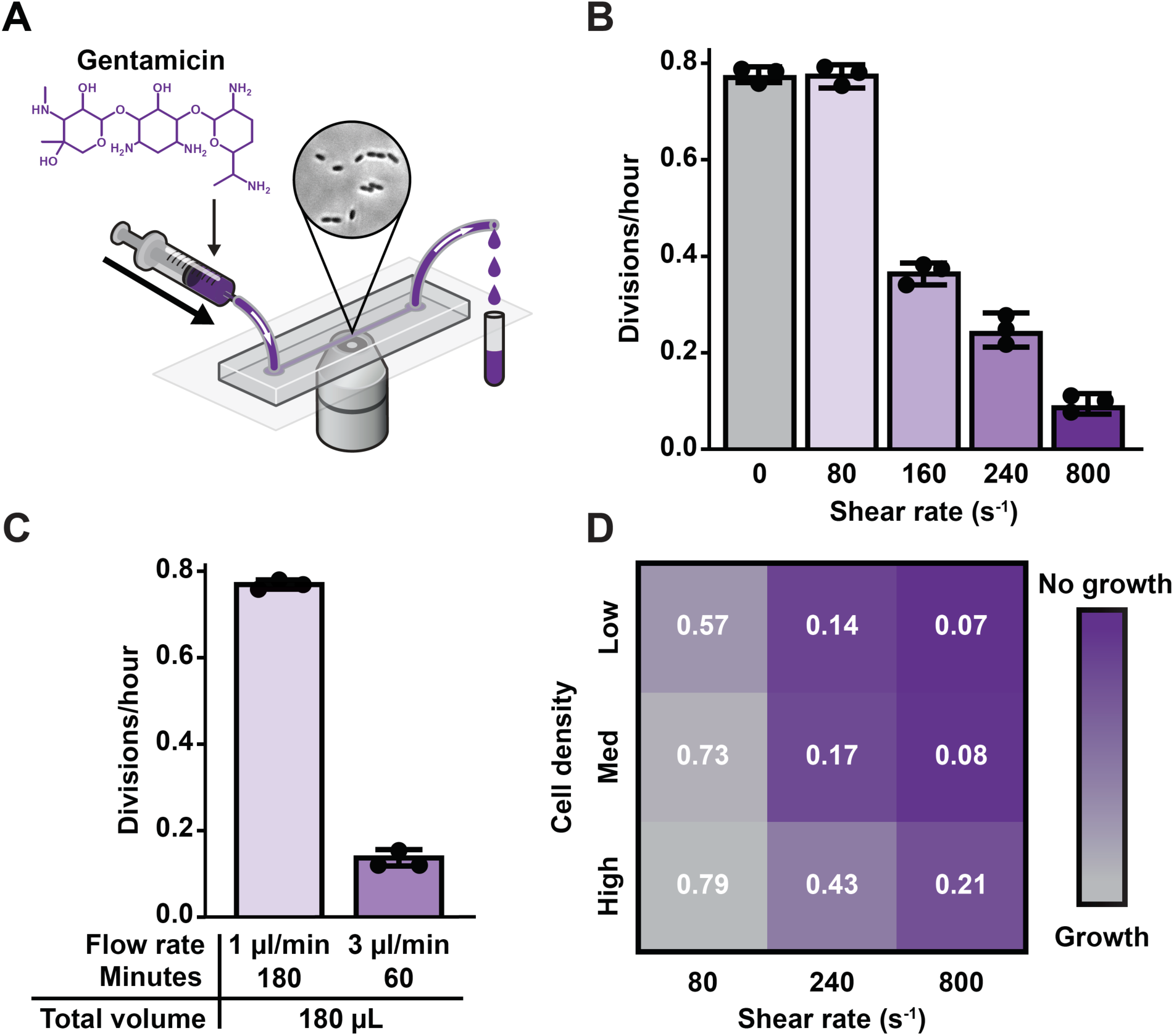
Shear rate and cell density collectively determine antibiotic effectiveness. (A) Illustration of the microfluidic setup used for exposing cells to medium supplemented with gentamicin. Channels are 50µm tall, 500 µm wide, and 2 cm long. Flow was generated with a syringe pump, and images were taken at the beginning of the channel. (B) Growth of gentamicin resistant P. aeruginosa cells at increasing shear rates over 4 hours in LB medium supplemented with 100 µg/mL gentamicin. (C) Experiments measuring gentamicin resistant cell growth with 100 µg/mL gentamicin at 1 µl/min for 180 minutes or 3 µl/min for 60 minutes. By changing both flow and duration, these experiments treated cells with equal total volumes of gentamicin. Experiment growth was measured in 60-minute bins, from 120-180 minutes for 1 µL/min and 0-60 minutes for 3 µL/min. (D) Checkerboard plot comparing gentamicin resistant cell growth across various shear rates and cell densities with 100 µg/mL gentamicin. Robust growth conditions are shown in grey which transition to purple for lesser growth conditions. Cell growth is imaged at the beginning of a 2-cm-long channel. For each biological replicate, 30 cells were chosen at random for quantification. Bar plots show SD error bars of 3 biological replicates, and checkerboard plot shows the average of three replicates.

How does flow overcome gentamicin resistance? By increasing flow in our experiments, we increased both the rate of antibiotic delivery and the total number of molecules delivered. To disentangle the effects of delivery and total dosage, we performed microfluidic experiments where we changed the flow rate while keeping the total dose constant. Specifically, we exposed *P. aeruginosa* cells to 180 µL of 100 µg/mL gentamicin in two different ways: first at a slower flow rate (1 µL/min) for a longer time (180 minutes) and second at a higher flow rate (3 µL/min) for a shorter time (60 minutes). We observed that cells exposed to the slower flow for a longer time grew robustly, while cells exposed to the higher flow for a shorter time did not (Figure 1C). Based on these results, we conclude that flow overcomes gentamicin resistance by promoting delivery independent of the total dose administered.

How does flow-driven delivery enhance gentamicin effectiveness? We hypothesized that gentamicin effectiveness is impacted by two rates: the rate of delivery by flow and the rate of inactivation by cells. To examine the combinatorial interactions between delivery and inactivation, we performed microfluidic experiments where we changed both flow (to alter delivery rate) and population density (to alter inactivation rate). Consistent with our hypothesis, increasing shear rate enhanced antibiotic effectiveness across all population densities, while increasing population density reduced antibiotic effectiveness across all shear rates (Figure 1D). Together, our microfluidic experiments support the hypothesis that shear rate enhances gentamicin effectiveness by outcompeting bacterial inactivation.

To quantitatively examine the interaction between shear rate and antibiotic inactivation, we simulated gentamicin delivery into populations with different cell densities. Our simulations included three key parameters: flow, diffusion, and gentamicin inactivation by cells. By holding flow and diffusion constant, we examined how the rate of gentamicin inactivation impacted delivery into a simulated population. At high cell density, inactivation was faster than delivery and gentamicin was eliminated from most of the channel (Figure 2A). At medium cell density, inactivation and delivery were more balanced and a spatial gradient of gentamicin was formed (Figure 2A). At low cell density, inactivation was slower than delivery and gentamicin levels remained high throughout the channel (Figure 2A). Thus, our simulations predict that inactivation rate can determine the spatial profile of gentamicin across bacterial populations.

**Figure 2:**
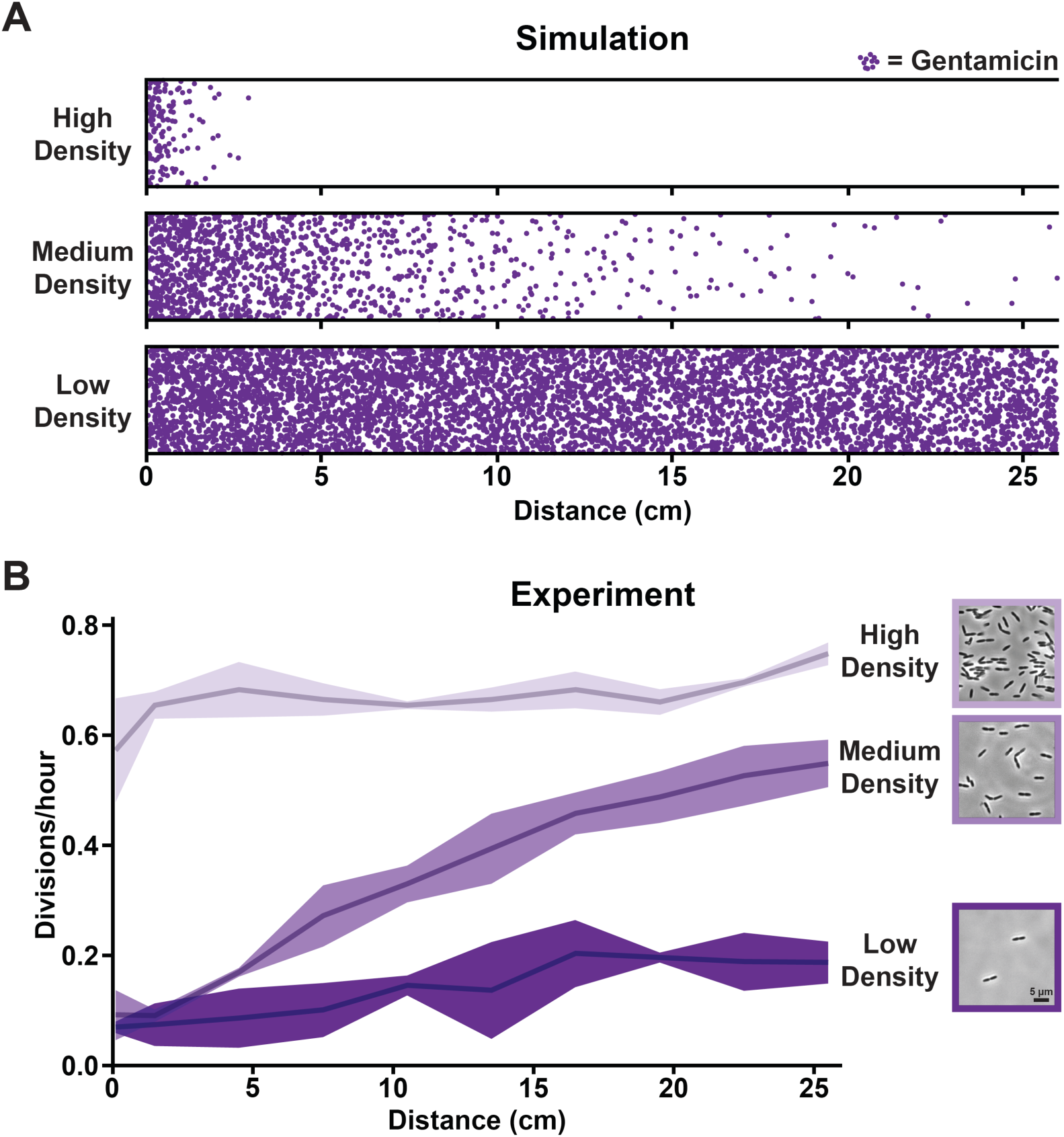
Bacterial antibiotic uptake patterns gradients of population growth. **(A)** Simulation of gentamicin molecules delivered in simulated microfluidic devices at 240 s^-1^ shear rate after 4 hours. Purple dots represent individual gentamicin molecules that experienced diffusion, flow from left to right, and removal when they contacted the bottom of the channel to represent bacterial removal. Bacterial removal rate of low density is 10-fold less than medium density, and removal rate for high density is 10-fold greater than medium density. **(B)** Experiments measuring gentamicin resistant *P. aeruginosa* population density growth for 4 hours with 100 µg/mL gentamicin at 240 s^-1^ shear rate. Channels are loaded from mid-log cultures (Medium Density) that are either diluted 10-fold (Low Density) or concentrated 10-fold (High Density). Cell density growth experiments in (B) closely matched the biophysical experiments in (A). Scale bar is 3 µm. For each biological replicate, 30 cells were chosen at random for quantification. Shaded regions show SD of three biological replicates.

To experimentally test our simulations, we used long microfluidic channels to study the spatial growth profiles of cells at different population densities. For these experiments, we seeded 27-cm-long channels (Figure S1) with cells at low, medium, and high density. We then exposed each population to 100 µg/mL gentamicin at a shear rate of 240 s^-1^. Supporting our simulations, a population at high cell density grew well throughout the channel and a population at low cell density grew poorly throughout the channel (Figure 2B). Further supporting our simulations, a medium cell density population grew poorly at the start of the channel and grew well at the end of the channel (Figure 2B). The resulting spatial gradients at medium density demonstrate that populations can defend themselves by inactivating gentamicin and flow can overwhelm this defense. Collectively, our results reveal that the balance between delivery and inactivation determines antibiotic effectiveness and shifts spatial population gradients.

Based on our spatial growth gradients, we hypothesized that bacterial populations can reduce the effective gentamicin concentration. To test our hypothesis, we treated a resistant *P. aeruginosa* population with gentamicin and used a bioassay to quantify the gentamicin concentration of the effluent (Figure 3A). First, we flowed 100 µg/mL gentamicin into a 27-cm-long channel with resistant cells and collected the effluent from the end of the channel. Second, we flowed the effluent into a second 2-cm-long channel with resistant cells and quantified cell growth. Third, we compared the resulting cell growth to a standard curve, which allowed us to estimate that the gentamicin concentration of the effluent was ∼18 µg/mL (Figure 3B). As the effluent had approximately 5-fold less gentamicin than at the start of the channel, we conclude that resistant *P. aeruginosa* populations can defend against gentamicin by reducing the effective concentration.

**Figure 3:**
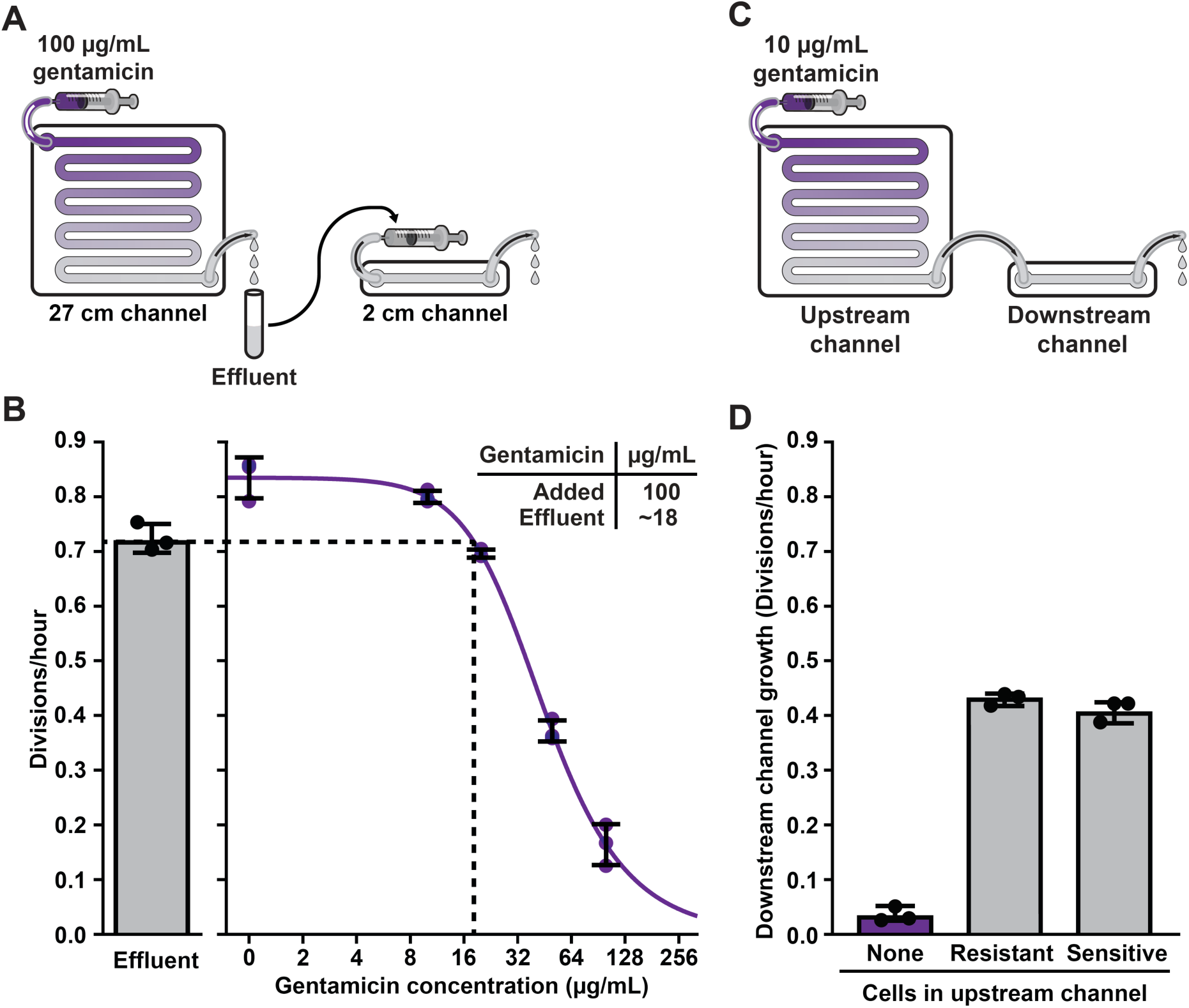
Resistant and sensitive bacteria both shape antibiotic gradients across populations. **(A)** Illustration of gentamicin bioassay setup for quantifying effluent concentration. Effluent was collected from 27 cm channels dosed with 100 µg/mL gentamicin at 240 s^-1^ shear rate. Then, effluent was flowed at 240 s^-1^ into 2 cm channels to measure cell growth. Both channels were loaded with gentamicin resistant *P. aeruginosa* cells. **(B)** Growth of gentamicin resistant cells in effluent matched with a standard growth curve of gentamicin. Regressing cell growth against the standard growth curve indicates gentamicin in filtered effluent was reduced to approximately 18 µg/mL. Standard was generated using a five-parameter logistic fit curve. **(C)** Illustration of dual microfluidic device setup for comparing population-level defense. Upstream channels used for gentamicin defense were dosed with 10 µg/mL gentamicin at 240 s^-1^ shear rate and were loaded with gentamicin resistant or sensitive cells. Downstream channels are used to measure effective concentration and were loaded with sensitive cells. Channels are connected via a 3-cm-long tubing bridge after loading with cells. **(D)** Growth of sensitive cells in the downstream channel after exposure to upstream channel effluent. Cell growth shows a similar level of reduced gentamicin concentrations from resistant and sensitive upstream populations. For each biological replicate, 30 cells were chosen at random for quantification. Error bars show SD of 3 biological replicates.

How do bacterial populations defend against gentamicin? As gentamicin likely enters cells through energy-dependent transporters that do not work in reverse (*39*, *40*), population-level defense could involve physical sequestration. However, as resistant cells chemically inactivate gentamicin, the roles of physical sequestration and chemical inactivation are unclear. To test the roles of sequestration and inactivation, we used a microfluidic approach to compare resistant populations (which can sequester and inactivate) to sensitive populations (which can only sequester). Our approach used two connected microfluidic channels (Figure 3C), where the first channel was the site of gentamicin defense and the second channel allowed us to measure the effective gentamicin concentration. Surprisingly, we observed that sensitive and resistant populations could equally lower the effective gentamicin concentration (Figure 3D), leading us to conclude that physical sequestration is very important to population-level defense.

Can populations of sensitive cells defend against gentamicin? As sensitive cells can lower the effective gentamicin concentration (Figure 3D), we hypothesized that sensitive populations would generate spatial growth gradients when treated with gentamicin. To test our hypothesis, we treated sensitive populations with a range of gentamicin concentrations (0-24 µg/mL) in a 27-cm-long microfluidic channel. Without gentamicin, cells grew robustly throughout the channel, and no growth gradient was observed (Figure 4A). At the highest gentamicin concentration (24 µg/mL), cells did not grow throughout the channel (Figure 4A). Supporting our hypothesis, all other gentamicin treatments (2, 3, 6, 12 µg/mL) resulted in spatial growth gradients with lower growth at the start of the channel and higher growth at the end of the channel (Figure 4A). In fact, the minimal inhibitory concentration for gentamicin at the start of the channel (6 µg/mL) was about 4-fold lower than the end of the channel (24 µg/mL) (Figure 4B). Thus, sensitive populations can defend against gentamicin, and this defense results in a situation where subpopulations of cells are differentially susceptible to antibiotic treatment.

**Figure 4:**
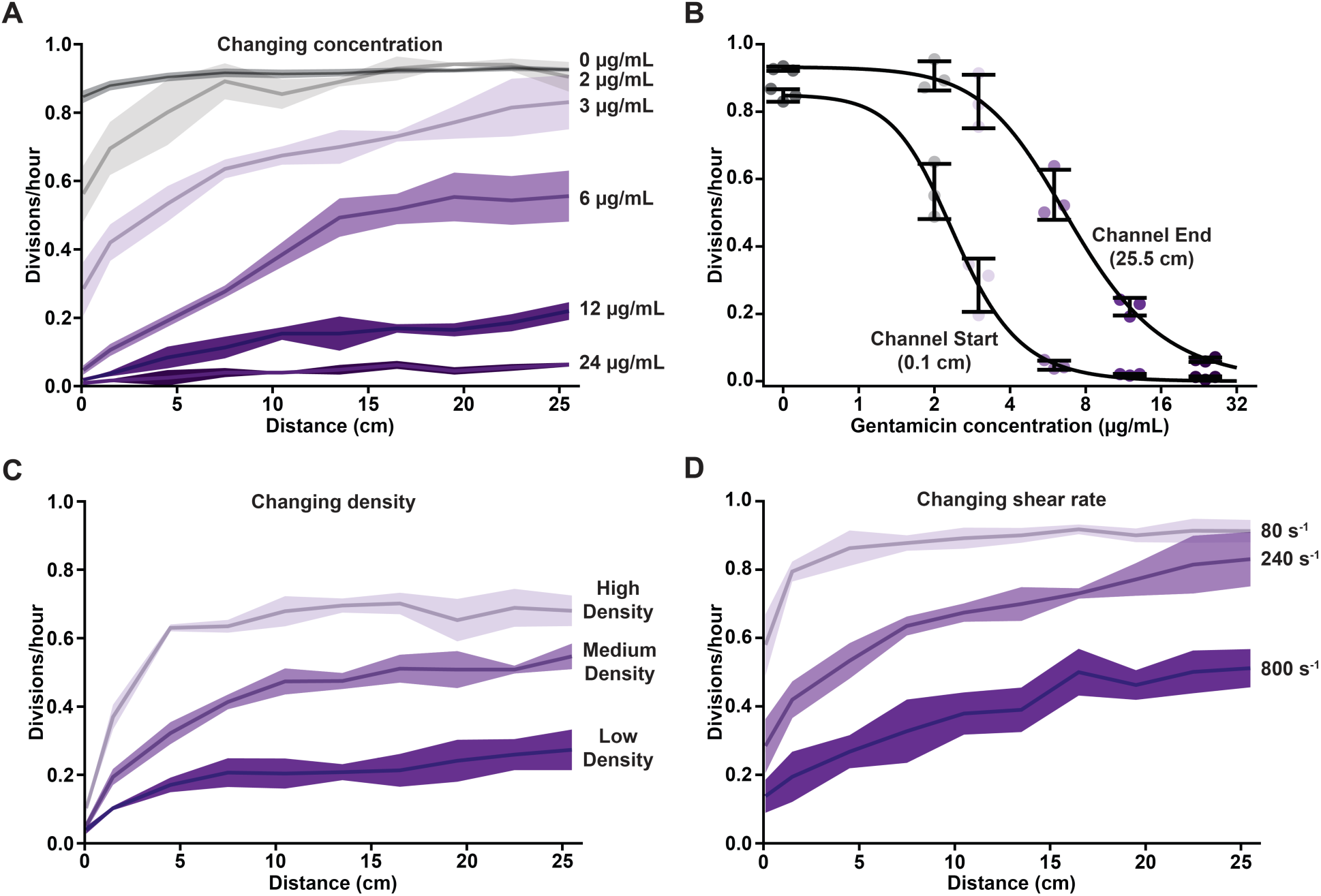
Shear rate and cell density impact antibiotic delivery into sensitive populations. **(A)** Experiments measuring gentamicin-sensitive *P. aeruginosa* population growth over 4 hours in increasing gentamicin concentrations at 240 s^-1^ shear rate and normal cell density. **(B)** Growth of *P. aeruginosa* cells at the start (0.1 cm) and end (25.5 cm) of the channel over increasing gentamicin concentrations. **(C)** Experiments measuring gentamicin-sensitive population growth over 4 hours in 6 µg/mL gentamicin at 240 s^-1^ shear rate at various population densities. Channels are loaded from mid-log cultures (Medium Density) that are either diluted 10-fold (Low Density) or concentrated 10-fold (High Density). **(D)** Experiments measuring gentamicin-sensitive population growth over 4 hours in 3 µg/mL gentamicin at medium density at various shear rates. Higher shear rates increase gentamicin effectiveness against sensitive populations. Gentamicin gradients developed have similar flat slopes to those gradients made with gentamicin-resistant populations (Figure 2B). All experiments were performed in 27-cm-long channels. Scale bar is 3 µm. For each biological replicate, 30 cells were chosen at random for quantification. Error bars and shaded regions show SD of 3 biological replicates.

Based on our new understanding, we hypothesized that the ability of sensitive populations to defend against gentamicin depends on the rates of antibiotic uptake and delivery. To test the role of gentamicin uptake, we altered population density in a 27-cm-long channel.

Specifically, we used a constant flow to deliver 6 µg/mL gentamicin into sensitive populations at low, medium, or high density. Consistent with our hypothesis that uptake is important, there was a stepwise increase in growth as population density increased (Figure 4C). To test the role of gentamicin delivery, we altered shear rate (80, 240, or 800 s^-1^) while holding population density constant. Consistent with our hypothesis that delivery is important, there was a stepwise decrease in growth as shear rate increased (Figure 4D). Based on these results, we conclude that uptake and delivery collectively determine gentamicin effectiveness across sensitive *P. aeruginosa* populations. Together, our microfluidic experiments demonstrate that flow enhances antibiotic effectiveness by delivery to overpower resistant and sensitive bacterial defenses.

## Discussion

Conventionally, antibiotics are discovered, developed, and tested in conditions lacking flow. However, during treatment, antibiotics are often delivered by flow. Our results demonstrate that flow has a major impact on antibiotic effectiveness, indicating that flow should be incorporated into antibiotic research and testing. Specifically, we discover that flow-based antibiotic delivery can overwhelm bacterial defenses, increasing penetration into bacterial populations and inhibiting bacterial growth. In non-flowing conditions (such as a Petri plate), cells in a population can defend themselves through chemical inactivation (Figure 5A) or physical sequestration (Figure 5B). Either way, cells at the interior are protected and can continue to grow. In flowing conditions (such as the bloodstream), our results suggest that flow-based delivery can overwhelm inactivation or sequestration (Figure 5C), thereby boosting antibiotic effectiveness and inhibiting bacterial growth. Our research highlights the limits of testing antibiotics with conventional assays and suggests that new approaches that incorporate flow are needed to precisely model the conditions bacteria face during infection.

**Figure 5:**
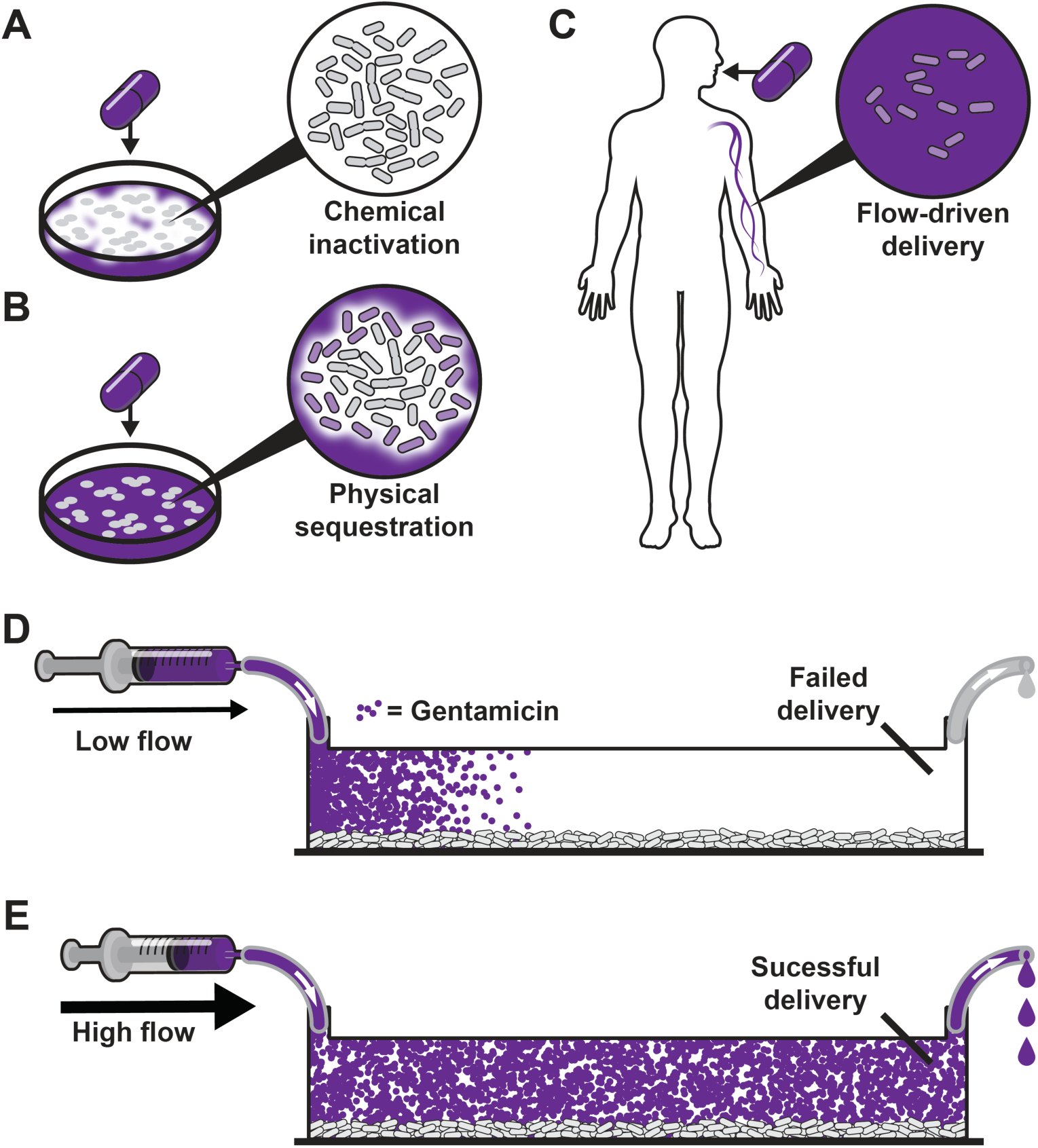
Flow-driven delivery boosts antibiotic effectiveness by overwhelming bacterial defenses. **(A)** On petri plates, resistant cells remove antibiotics by chemical inactivation, completely clearing antibiotic and can grow. **(B)** On petri plates, sensitive cells remove antibiotics by physical sequestration, where exterior cells generate local zones of clearance so only interior cells can grow. **(C)** In host-relevant conditions, resistant or sensitive cells experience flow, cannot remove antimicrobials quickly enough, and do not grow. **(D)** In low flow, bacterial populations remove antibiotics more quickly than they are delivered, allowing for growth and protecting downstream cells. **(E)** In high flow, gentamicin is rapidly replenished more than it is removed by cells and overwhelms the entire bacterial population.

How does flow boost antibiotic effectiveness? Collectively, our data support a model where flow boosts antibiotic effectiveness by delivering antibiotics faster than bacterial populations can remove them. By changing shear rate (Figure 1B), we showed that increasing flow increases gentamicin effectiveness. By changing population density (Figures 1D and 2B), we showed that increasing removal decreases gentamicin effectiveness. Using a combinatorial approach, we demonstrated that the balance between delivery and removal ultimately sets the effectiveness of gentamicin (Figure 1D). To address an alternate hypothesis that flow boosts effectiveness by delivering more antibiotic, we designed an experiment where flow was changed while the total dose delivered was held constant (Figure 1C). This experiment conclusively showed that the effect of flow does not depend on an increased total dose (Figure 1C), further supporting our model. Thus, we conclude that flow boosts antibiotic effectiveness by delivering antibiotics faster than bacterial populations can remove them.

Gentamicin is an aminoglycoside antibiotic that gets into cells and targets protein synthesis. Cells can protect themselves against gentamicin by chemical inactivation via an acetyltransferase (Figure S2). Our results indicate that cell populations can also protect against gentamicin by physical sequestration. There are multiple hypotheses to explain physical sequestration of gentamicin: adsorption to the channel, adsorption to the cell surface, or internalization within cells. As our empty channel control (Figure 3D) did not reduce the effective gentamicin concentration, we refute the channel adsorption hypothesis. As cell density is important (Figure 4C), we think that both cell surface adsorption hypothesis and internalization hypothesis are reasonable. We prefer the internalization hypothesis, due to the fact that gentamicin is entering cells and inhibiting growth. As aminoglycosides enter cells through energy-dependent processes (*39–44*), it is likely that gentamicin gets into cells and does not get out.

By using our microfluidic platform to examine the interactions between biology (*P. aeruginosa*), chemistry (gentamicin), and physics (shear flow), we have provided a new framework to better understand antibiotic effectiveness in dynamic environments. Specifically, we have learned how flow can enhance gentamicin effectiveness by overwhelming *P. aeruginosa* defense strategies. Our experiments revealed that sensitive *P. aeruginosa* cells can protect themselves through population-level physical sequestration, generating a subpopulation of cells that avoid treatment and continue to grow. Our results highlight the need to study antibiotics in flowing conditions and should inspire the use of flow in future efforts to more precisely test antibiotic effectiveness.

## Acknowledgments

We thank Anuradha Sharma, Piyush Sharma, Evan Johnson, Iota Chen, Mickey Marcheschi, Jim Imlay, Wilfred van der Donk, and Zemer Gitai for helpful discussions and comments on the manuscript.

## Funding

This work was supported by grant R35GM155443 from the National Institutes of Health to J.E.S. and grant R35GM155280 from the National Institutes of Health to M.D.K. This work was also supported by grant #25-5910 from the Roy J. Carver Charitable Trust to J.E.S.

## Contributions

A.M.S., M.D.K., and J.E.S. designed research. A.M.S. and M.D.K. performed research. A.M.S., M.D.K., and J.E.S. analyzed data. A.M.S. and J.E.S. wrote the paper.

## Supplementary Information for

### Materials and Methods

#### Bacterial strains and growth conditions

*Pseudomonas aeruginosa* strains used in this paper are PA14 Δ*pilA*::FRT (JS176) and PA14 Δ*pilA*::*aacC1* (JS177), which provides resistance to gentamicin. *P. aeruginosa* cultures were grown in LB broth on a cell culture roller drum, and on LB plates (1.5% Bacto Agar) at 37°C. LB broth was prepared using LB broth Miller (BD Biosciences).

#### Preparation of antimicrobial solutions

LB with gentamicin was generated as previously described (*35*). Briefly, stock solutions were prepared from gentamicin sulfate (GoldBio) dissolved in sterile water. Final gentamicin concentrations were diluted in LB medium to the desired concentration.

#### Fabrication of microfluidic devices

Microfluidic devices were prepared as previously described (*24*). Briefly, photomasks were designed on Illustrator (Adobe Creative Suite) and printed by Artnet Pro, Inc. Using soft lithography techniques, photomask patterns were transferred onto 100 mm silicon wafers (University Wafer) that were spin coated with SU-8 3050 photoresist (MicroChem). Microfluidic devices are made of Polydimethylsiloxane (Dow SYLGARD 184 Kit) at a 1:10 ratio and plasma-treated to bond on a 60 mm x 35 mm x 0.16 mm superslip micro cover glass (Ted Pella, Inc.). Growth experiments used channels with dimensions of 500 µm wide x 50 µm tall x 2 cm long or 500 µm wide x 50 µm tall x 27 cm long with 8 turns.

#### Phase contrast microscopy

Timelapse images were captured on a Nikon ECLIPSE Ti2-E inverted microscope using the NIS Elements interface. For all imaging, we used a Nikon 40x Plan Ph2 0.95 NA objective, a Hamamatsu ORCA-Flash 4.0 LT3 Digital CMOS camera, and a Lumencor SOLA Light Engine LED light source.

#### Preparation of microfluidic devices with bacteria

Microfluidic devices were loaded with bacteria as previously described (*24*). Briefly, all experiments were performed at approximately 22°C and with mid-log bacterial cultures. Cultures were loaded into microfluidic devices via a syringe (BD Biosciences) on a syringe pump (KD Scientific Legato 210) set at 10 µL/min. Cells were allowed to adhere for 10 minutes before experimental flow exposure. For experiments changing population density, mid-log bacterial cultures were either diluted or concentrated 10-fold in LB medium prior to loading into microfluidic devices. Device inlets are attached to a length of polyethylene tubing, ID 0.015” x OD 0.043” (Brain Tree Scientific) that is sheathed over a 26-gauge x 1/2” hypodermic needle (Air-tite Products) that is affixed on a syringe. Device outlets contain another length of tubing for vacating spend medium into a bleach-containing waste container. Medium-filled syringes were fastened on a syringe pump that was set at flow rates between 1 - 10 µl/min, which corresponds with shear rates between 80 - 800 s^-1^. For microfluidic effluent filtration experiments, effluent was collected over 4 hours at a flow rate of 3 µl/min and filtered through a 0.22 µm size syringe filter (MilliporeSigma) with a syringe. For direct microfluidic effluent exposure experiments, mid-log cultures were loaded separately via syringe pump then connected via a short tubing bridge approximately 3 cm long.

#### Shear rate calculations

Shear rate experienced in microfluidic devices was calculated using this equation:

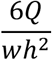

Where *Q* is flow rate, *w* is channel width, and *h* is channel height. Shear rate is specified throughout in units of s^-1^.

#### Quantification of cellular growth

Cell growth measurements were performed as previously described (*35*). Briefly, measuring cell growth during gentamicin treatments with and without flow, images were captured every 10 minutes for up to 4 hours. Cell growth quantification was performed manually on ImageJ software by counting total divisions and dividing by time. At least 30 cells per field were chosen at random for growth tracking. All growth experiments were performed with cells lacking *pilA*, which prevented twitching motility and allowed for easy cell tracking.

#### Biophysical simulations

Simulations of molecular advection and diffusion were performed as previously described (*24*). Briefly, simulated channels were randomly filled with molecules that were assigned diffusive behavior by combining laminar transport and Brownian dynamic simulations. Channel parabolic flow speed profiles were modeled using the Hagen-Poiseuille equation. Molecules were allowed to diffuse along the x-axis and y-axis of the channel. Gentamicin was simulated with a diffusion coefficient of 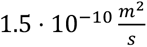. Molecular removal was changed to simulate cell density in channels.

Normal density was estimated to be 1 of every 100 molecules that reached the channel bottom (where cells are located). For high and low density, removal was increased or decreased by 10-fold (1:10 for high density and 1:1000 for low density).

---

**Figure S1:**
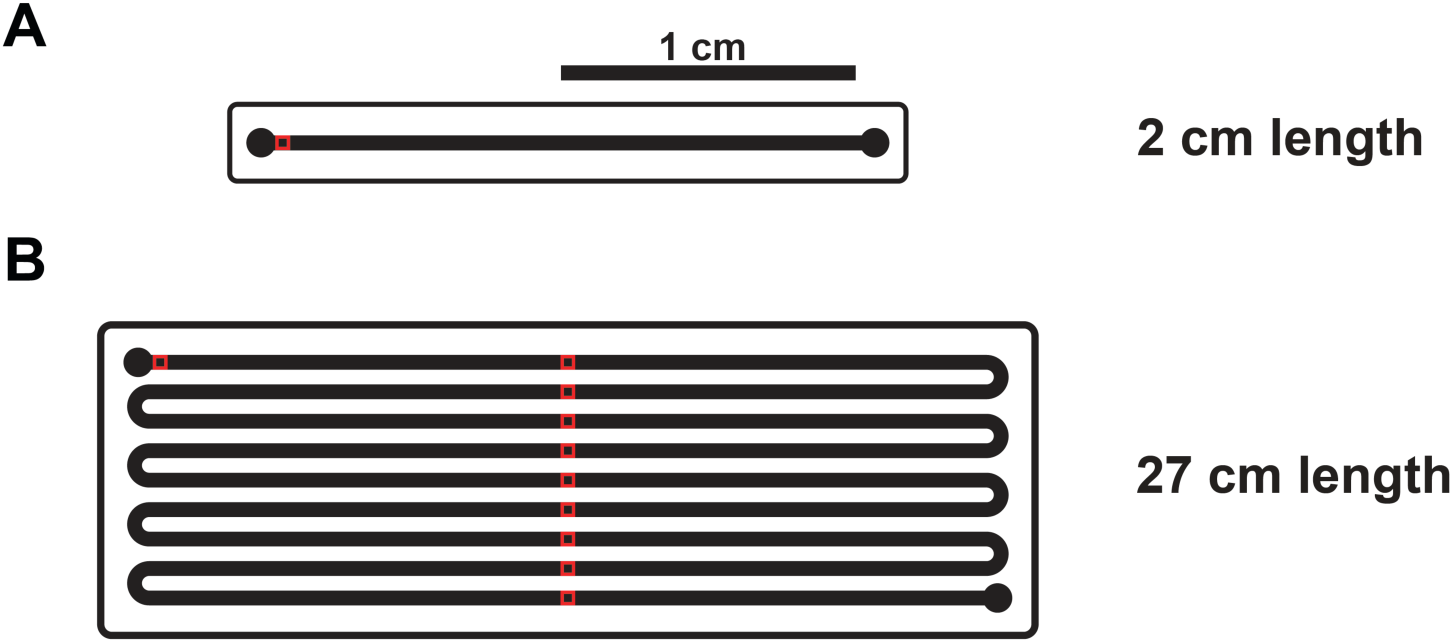
Designs of microfluidic devices used in this study. **(A)** Top-down view of a 2-cm-long device. **(B)** Top-down view of a 27-cm-long device containing 8 turns. Scale bar is 1 cm. Small red boxes in channel designs depict the approximate regions that were used for imaging. Specifically, 2-cm-long devices were imaged at the beginning of the channel (0.1 cm). 27-cm-long devices were imaged at 0.1, 1.5, 3.5, 7.5, 10.5, 13.5, 16.5, 19.5, 22.5, and 25.5 cm distances. For 2 cm channels, 6 independent experiments can be performed simultaneously. For 27 cm channels, 3 independent experiments can be performed simultaneously.

**Figure S2:**
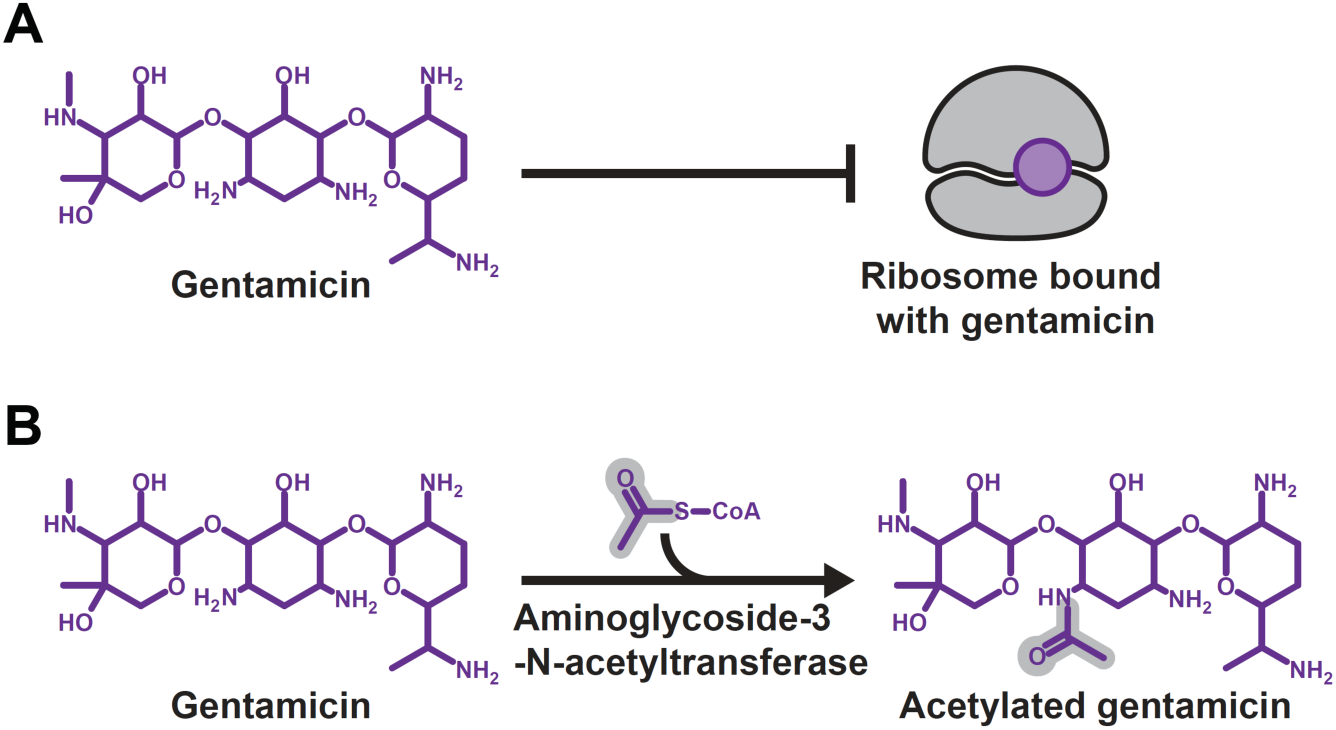
Gentamicin mechanism of action and resistance in this study. **(A)** Gentamicin binds to ribosomes to inhibit protein translation (*45*). **(B)** Gentamicin resistant cells modify gentamicin by encoding an aminoglycoside-3-N-acetyltransferase that transfers the acetyl group from acetyl-CoA to the 3C amino group of gentamicin (*39*).

**Figure S3:**
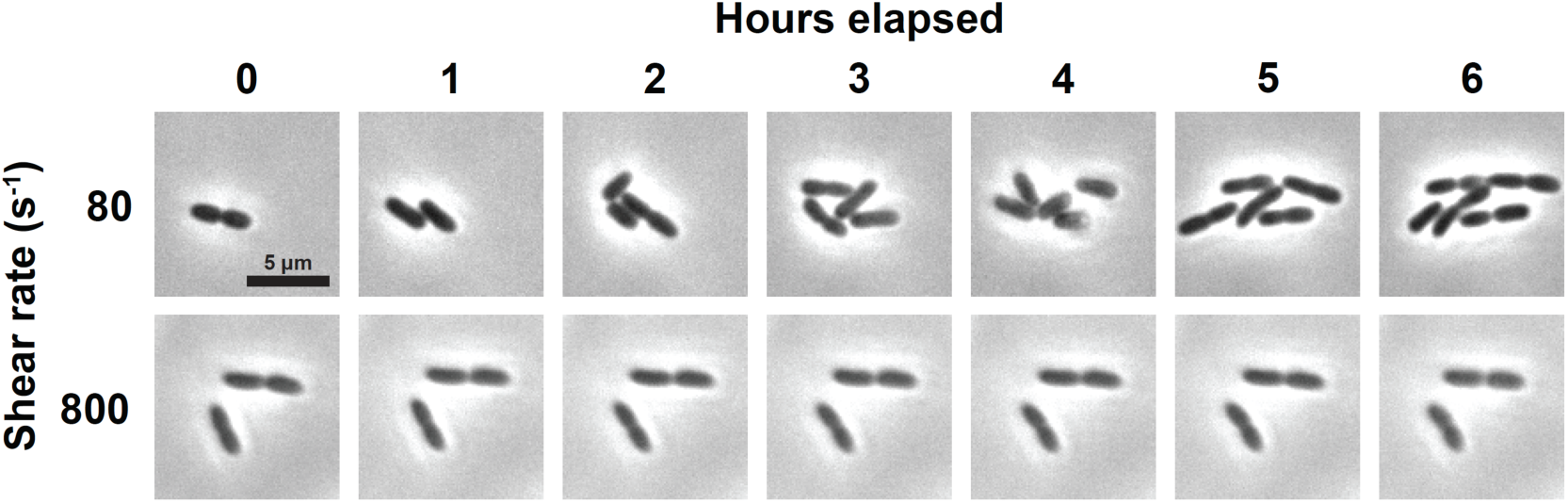
Representative images of shear rate increasing gentamicin effectiveness. Phase contrast images representative of three biological replicates of gentamicin resistant *P. aeruginosa* Δ*pilA* cells at various shear rates over 6 hours. Images were taken at the beginning (∼0.1cm) of a 2-cm-long microfluidic channel. Cells are exposed to either 80 s^-1^ or 800 s^-1^ shear rates in LB medium supplemented with 100 µg/mL gentamicin. Scale bar is 5 µm.

**Figure S4:**
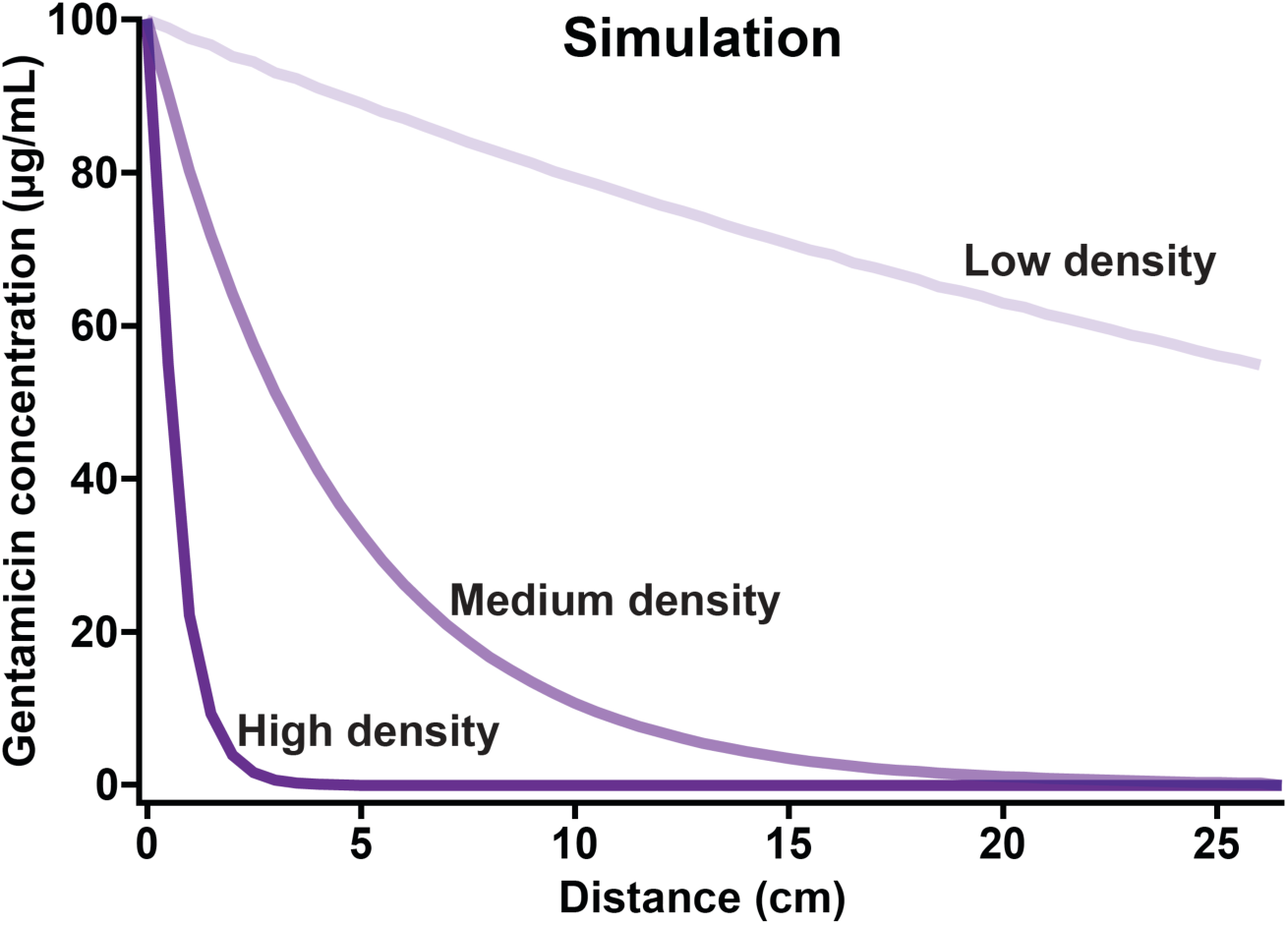
Biophysical simulations predict that increasing cell density shortens spatial gradients. Quantification of gentamicin concentrations across the length of a simulated 27-cm-long microfluidic channels at various population densities.

**Figure S5:**
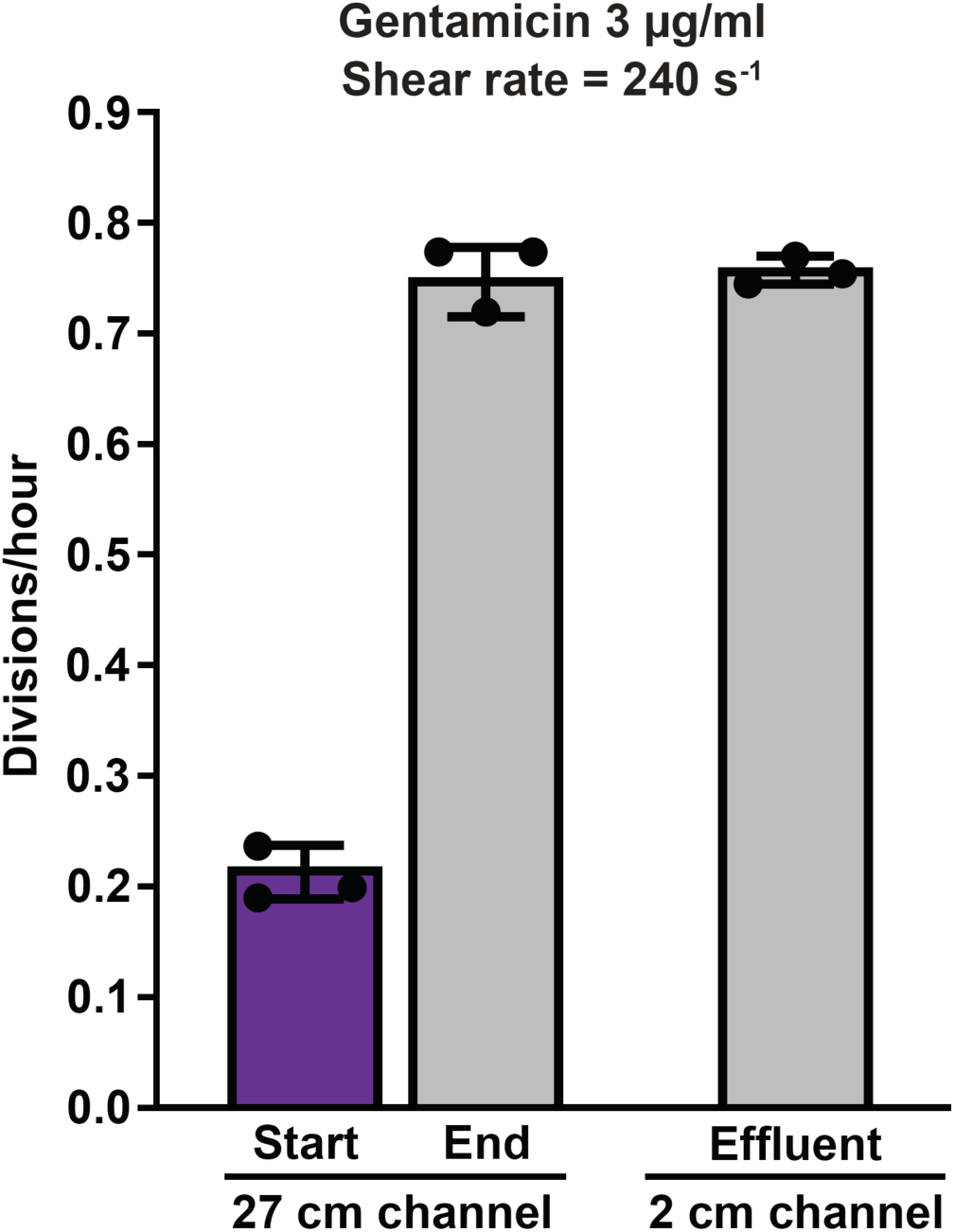
Sensitive cells remove gentamicin from filtered effluent. Growth of gentamicin sensitive *P. aeruginosa* cells across a 27-cm-long channel and in filtered effluent. Upstream channels were dosed with LB medium supplemented with 3 µg/mL gentamicin and upstream and downstream cells were exposed at 240 s^-1^ shear rate. Error bars show SD of 3 biological replicates.

**Figure S6:**
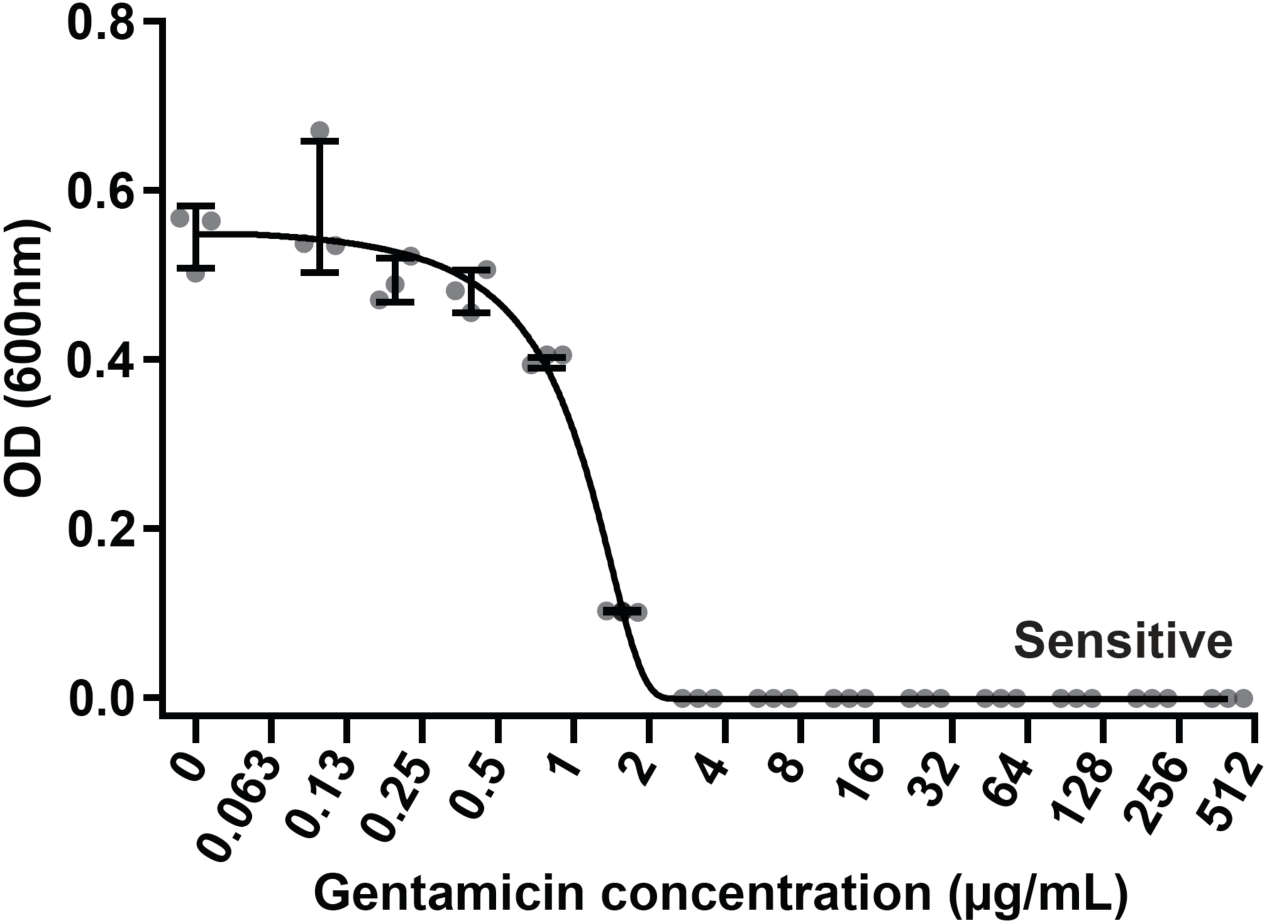
Gentamicin sensitive cell growth curve. Sensitive *P. aeruginosa* cell growth over increasing gentamicin concentrations. Growth was measured after 8 hours. Error bars show SD of 3 biological replicates.

**Table S1:** ***P. aeruginosa* strains used in this study**

| ***P. aeruginosa* strains** | **Description** | **Source** |
| --- | --- | --- |
| PA14 | Wildtype; clinical isolate from burn wound | (46) |
| JS176 | PA14 $\Delta pilA::FRT$ | (5) |
| JS177 | PA14 $\Delta pilA::aacC1$ | (5) |

## Notes

### Competing Interest Statement

The authors have declared no competing interest.

## References

1 G. C. Padron, A. M. Shuppara, J.-J. S. Palalay, A. Sharma, J. E. Sanfilippo, Bacteria in Fluid Flow. J. Bacteriol. 205, e00400–22 (2023).

2 A. Persat, C. D. Nadell, M. K. Kim, F. Ingremeau, A. Siryaporn, K. Drescher, N. S. Wingreen, B. L. Bassler, Z. Gitai, H. A. Stone, The Mechanical World of Bacteria. Cell 161, 988–997 (2015).

3 J. D. Wheeler, E. Secchi, R. Rusconi, R. Stocker, Not Just Going with the Flow: The Effects of Fluid Flow on Bacteria and Plankton. Annu. Rev. Cell Dev. Biol. 35, 213–237 (2019).

4. J.-J. S. Palalay, J. E. Sanfilippo, Flow-induced bending of flagella restricts Pseudomonas aeruginosa surface departure. mBio 17, e0274025 (2026).

5. J.-J. S. Palalay, A. N. Simsek, J. L. Reed, M. D. Koch, B. Sabass, J. E. Sanfilippo, Shear force enhances adhesion of Pseudomonas aeruginosa by counteracting pilus-driven surface departure. Proc. Natl. Acad. Sci. 120, e2307718120 (2023).

6 S. Lecuyer, R. Rusconi, Y. Shen, A. Forsyth, H. Vlamakis, R. Kolter, H. A. Stone, Shear Stress Increases the Residence Time of Adhesion of *Pseudomonas aeruginosa*. Biophys. J. 100, 341–350 (2011).

7 E. Secchi, A. Vitale, G. L. Miño, V. Kantsler, L. Eberl, R. Rusconi, R. Stocker, The effect of flow on swimming bacteria controls the initial colonization of curved surfaces. Nat. Commun. 11, 2851 (2020).

8 K. M. Hallinen, S. P. Bodine, H. A. Stone, T. W. Muir, N. S. Wingreen, Z. Gitai, Bacterial species with different nanocolony morphologies have distinct flow-dependent colonization behaviors. Proc. Natl. Acad. Sci. 122, e2419899122 (2025).

9 A. Siryaporn, M. K. Kim, Y. Shen, H. A. Stone, Z. Gitai, Colonization, Competition, and Dispersal of Pathogens in Fluid Flow Networks. Curr. Biol. 25, 1201–1207 (2015).

10 Y. Shen, A. Siryaporn, S. Lecuyer, Z. Gitai, H. A. Stone, Flow Directs Surface-Attached Bacteria to Twitch Upstream. Biophys. J. 103, 146–151 (2012).

11 R. Tao, A. Théry, S. Que, A. J. T. M. Mathijssen, Invasion of bacteria swimming upstream into microstructured devices. Newton 2 (2026).

12 A. Kannan, Z. Yang, M. K. Kim, H. A. Stone, A. Siryaporn, Dynamic switching enables efficient bacterial colonization in flow. Proc. Natl. Acad. Sci. 115, 5438–5443 (2018).

13 T. Kaya, H. Koser, Direct Upstream Motility in *Escherichia coli*. Biophys. J. 102, 1514–1523 (2012).

14 R. Rusconi, J. S. Guasto, R. Stocker, Bacterial transport suppressed by fluid shear. Nat. Phys. 10, 212–217 (2014).

15 A. J. T. M. Mathijssen, N. Figueroa-Morales, G. Junot, É. Clément, A. Lindner, A. Zöttl, Oscillatory surface rheotaxis of swimming *E. coli* bacteria. Nat. Commun. 10, 3434 (2019).

16 K. Drescher, Y. Shen, B. L. Bassler, H. A. Stone, Biofilm streamers cause catastrophic disruption of flow with consequences for environmental and medical systems. Proc. Natl. Acad. Sci. 110, 4345–4350 (2013).

17 R. Rusconi, S. Lecuyer, L. Guglielmini, H. A. Stone, Laminar flow around corners triggers the formation of biofilm streamers. J. R. Soc. Interface 7, 1293–1299 (2010).

18 R. Rusconi, S. Lecuyer, N. Autrusson, L. Guglielmini, H. A. Stone, Secondary Flow as a Mechanism for the Formation of Biofilm Streamers. Biophys. J. 100, 1392–1399 (2011).

19 G. Savorana, T. Redaelli, D. Truzzolillo, L. Cipelletti, E. Secchi, Stress-hardening behaviour of biofilm streamers. Nat. Commun. 16, 9497 (2025).

20 G. C. Padron, S. Chen, A. Sharma, Z. Modi, M. D. Koch, J. E. Sanfilippo, Shear flow promotes bacterial growth and shapes spatial gradients by rapidly replenishing scarce nutrients. mBio 17, e03446–25 (2026).

21 J. P. H. Wong, M. Fischer-Stettler, S. C. Zeeman, T. J. Battin, A. Persat, Fluid flow structures gut microbiota biofilm communities by distributing public goods. Proc. Natl. Acad. Sci. 120, e2217577120 (2023).

22 K. Drescher, C. D. Nadell, H. A. Stone, N. S. Wingreen, B. L. Bassler, Solutions to the Public Goods Dilemma in Bacterial Biofilms. Curr. Biol. 24, 50–55 (2014).

23 J. Nguyen, V. Fernandez, S. Pontrelli, U. Sauer, M. Ackermann, R. Stocker, A distinct growth physiology enhances bacterial growth under rapid nutrient fluctuations. Nat. Commun. 12, 3662 (2021).

24 G. C. Padron, A. M. Shuppara, A. Sharma, M. D. Koch, J.-J. S. Palalay, J. N. Radin, T. E. Kehl-Fie, J. A. Imlay, J. E. Sanfilippo, Shear rate sensitizes bacterial pathogens to H_2_O_2_ stress. Proc. Natl. Acad. Sci. 120, e2216774120 (2023).

25 A. Sharma, A. M. Shuppara, G. C. Padron, J. E. Sanfilippo, Combining multiple stressors blocks bacterial migration and growth. Curr. Biol. 34, 5774–5781.e4 (2024).

26 J. E. Sanfilippo, A. Lorestani, M. D. Koch, B. P. Bratton, A. Siryaporn, H. A. Stone, Z. Gitai, Microfluidic-based transcriptomics reveal force-independent bacterial rheosensing. Nat. Microbiol. 4, 1274–1281 (2019).

27 M. K. Kim, F. Ingremeau, A. Zhao, B. L. Bassler, H. A. Stone, Local and global consequences of flow on bacterial quorum sensing. Nat. Microbiol. 1, 1–5 (2016).

28 M. J. Kirisits, J. J. Margolis, B. L. Purevdorj-Gage, B. Vaughan, D. L. Chopp, P. Stoodley, M. R. Parsek, Influence of the Hydrodynamic Environment on Quorum Sensing in *Pseudomonas aeruginosa* Biofilms. J. Bacteriol. 189, 8357–8360 (2007).

29 K. Lewis, Recover the lost art of drug discovery. Nature 485, 439–440 (2012).

30 M. A. Cook, G. D. Wright, The past, present, and future of antibiotics. Sci. Transl. Med. 14, eabo7793 (2022).

31 T. Bjarnsholt, M. Whiteley, K. P. Rumbaugh, P. S. Stewart, P. Ø. Jensen, N. Frimodt-Møller, The importance of understanding the infectious microenvironment. Lancet Infect. Dis. 22, 88–92 (2022).

32 E. M. Darby, E. Trampari, P. Siasat, M. S. Gaya, I. Alav, M. A. Webber, J. M. A. Blair, Molecular mechanisms of antibiotic resistance revisited. Nat. Rev. Microbiol. 21, 280–295 (2023).

33. J. Prince, A.-A. D. Jones, Heterogenous biofilm mass-transport model replicates periphery sequestration of antibiotics in *Pseudomonas aeruginosa* PAO1 microcolonies. *Proc. Natl. Acad. Sci*. 120, e2312995120 (2023).

34 B. S. Tseng, W. Zhang, J. J. Harrison, T. P. Quach, J. L. Song, J. Penterman, P. K. Singh, D. L. Chopp, A. I. Packman, M. R. Parsek, The extracellular matrix protects *Pseudomonas aeruginosa* biofilms by limiting the penetration of tobramycin. Environ. Microbiol. 15, 2865–2878 (2013).

35 A. M. Shuppara, G. C. Padron, A. Sharma, Z. Modi, M. D. Koch, J. E. Sanfilippo, Shear flow patterns antimicrobial gradients across bacterial populations. Sci. Adv. 11, eads5005 (2025).

36 J. J. Paszkowiak, A. Dardik, Arterial Wall Shear Stress: Observations from the Bench to the Bedside. Vasc. Endovascular Surg. 37, 47–57 (2003).

37 Y.-H. Lee, C.-W. Lai, Y.-C. Cheng, Fluid Shear Stress Induces Cell Cycle Arrest in Human Urinary Bladder Transitional Cell Carcinoma Through Bone Morphogenetic Protein Receptor-Smad1/5 Pathway. Cell. Mol. Bioeng. 11, 185–195 (2018).

38 W. Salibe-Filho, T. L. S. Araujo, E. G. Melo, L. B. C. T. Coimbra, M. S. Lapa, M. M. P. Acencio, O. Freitas-Filho, V. L. Capelozzi, L. R. Teixeira, C. J. C. S. Fernandes, F. B. Jatene, F. R. M. Laurindo, M. Terra-Filho, Shear stress-exposed pulmonary artery endothelial cells fail to upregulate HSP70 in chronic thromboembolic pulmonary hypertension. PLOS ONE 15, e0242960 (2020).

39 M. Lang, A. Carvalho, Z. Baharoglu, D. Mazel, Aminoglycoside uptake, stress, and potentiation in Gram-negative bacteria: new therapies with old molecules. Microbiol. Mol. Biol. Rev. 87, e0003622 (2023).

40 W. W. Nichols, S. N. Young, Respiration-dependent uptake of dihydrostreptomycin by *Escherichia coli*. Its irreversible nature and lack of evidence for a uniport process. Biochem. J. 228, 505–512 (1985).

41 K. Andry, R. C. Bockrath, Dihydrostreptomycin accumulation in *E. coli*. Nature 251, 534–536 (1974).

42 S. M. Mates, L. Patel, H. R. Kaback, M. H. Miller, Membrane potential in anaerobically growing *Staphylococcus aureus* and its relationship to gentamicin uptake. Antimicrob. Agents Chemother. 23, 526–530 (1983).

43 P. D. Damper, W. Epstein, Role of the membrane potential in bacterial resistance to aminoglycoside antibiotics. Antimicrob. Agents Chemother. 20, 803–808 (1981).

44 H. S. Fraimow, J. B. Greenman, I. M. Leviton, T. J. Dougherty, M. H. Miller, Tobramycin uptake in *Escherichia coli* is driven by either electrical potential or ATP. J. Bacteriol. 173, 2800–2808 (1991).

45 K. M. Krause, A. W. Serio, T. R. Kane, L. E. Connolly, Aminoglycosides: An Overview. *Cold Spring Harb. Perspect. Med*. 6, a027029 (2016).

46 L. G. Rahme, E. J. Stevens, S. F. Wolfort, J. Shao, R. G. Tompkins, F. M. Ausubel, Common Virulence Factors for Bacterial Pathogenicity in Plants and Animals. Science 268, 1899–1902 (1995).

